# Multiple phage resistance systems inhibit infection via SIR2-dependent NAD^+^ depletion

**DOI:** 10.1101/2021.12.14.472415

**Authors:** Jeremy Garb, Anna Lopatina, Aude Bernheim, Mindaugas Zaremba, Virginijus Siksnys, Sarah Melamed, Azita Leavitt, Adi Millman, Gil Amitai, Rotem Sorek

**Author notes:** Division of Microbial Ecology, Department of Microbiology and Ecosystem Science, Center for Microbiology and Environmental Systems Science, University of Vienna, Vienna, Austria (Anna Lopatina). Université de Paris, IAME, UMR 1137, INSERM, Paris, France.

## Abstract

Defense-associated sirtuins (DSR) comprise a family of proteins that defend bacteria from phage infection via an unknown mechanism. These proteins are common in bacteria and harbor an N-terminal sirtuin (SIR2) domain. In this study we report that DSR proteins degrade nicotinamide adenine dinucleotide (NAD^+^) during infection, depleting the cell of this essential molecule and aborting phage propagation. Our data show that one of these proteins, DSR2, directly identifies phage tail tube proteins and then becomes an active NADase in *Bacillus subtilis*. Using a phage mating methodology that promotes genetic exchange between pairs of DSR2-sensitive and DSR2–resistant phages, we further show that some phages express anti-DSR2 proteins that bind and repress DSR2. Finally, we demonstrate that the SIR2 domain serves as an effector NADase in a diverse set of phage defense systems outside the DSR family. Our results establish the general role of SIR2 domains in bacterial immunity against phages.

## Main text

SIR2-domain proteins, or sirtuins, are found in organisms ranging from bacteria to humans. These proteins have been widely studied in yeast and mammals, where they were shown to regulate transcription repression, recombination, DNA repair and cell cycle processes (North and Verdin, 2004). In eukaryotes, SIR2 domains were shown to possess enzymatic activities, and function either as protein deacetylases or ADP ribosyltransferases (Imai et al., 2000; Tanny et al., 1999). In both cases, the SIR2 domain utilizes nicotinamide adenine dinucleotide (NAD^+^) as a cofactor for the enzymatic reaction (Dang and Pfizer, 2014).

In bacteria, SIR2 domains were recently shown to participate in defense systems that protect against phages. These domains are associated with multiple different defense systems, including prokaryotic argonautes (pAgo) (Makarova et al., 2009), Thoeris (Doron et al., 2018; Ofir et al., 2021), AVAST, DSR, and additional systems (Gao et al., 2020). It was recently shown that the SIR2 domain in the Thoeris defense system is an NADase responsible for depleting NAD^+^ from the cell once phage infection has been sensed (Ofir et al., 2021). However, it is currently unknown whether SIR2 domains in other defense systems perform a similar function, or whether they have other roles in phage defense.

To study the role of SIR2 domains in bacterial anti-phage defense, we began by focusing on DSR2, a minimal defense system that includes a single protein with an N-terminal SIR2 domain and no additional identifiable domains (Figure 1A). The DSR2 gene family was recently identified based on a screen for genes commonly found in bacterial anti-phage defense islands (Gao et al., 2020). We cloned the DSR2 gene from *Bacillus subtilis* 29R, under the control of its native promoter, into the genome of *B. subtilis* BEST7003 which naturally lacks this gene, and challenged the DSR2-containing strain with a set of phages from the SPbeta family. We found that DSR2 protected *B. subtilis* against phage SPR (Figure 1B). Point mutations in residues N133 and H171 in DSR2, both of which predicted to disable the active site of the SIR2 domain, abolished defense, suggesting that the enzymatic activity of SIR2 is essential for DSR2 defense (Figure 1B).

**Figure 1.**
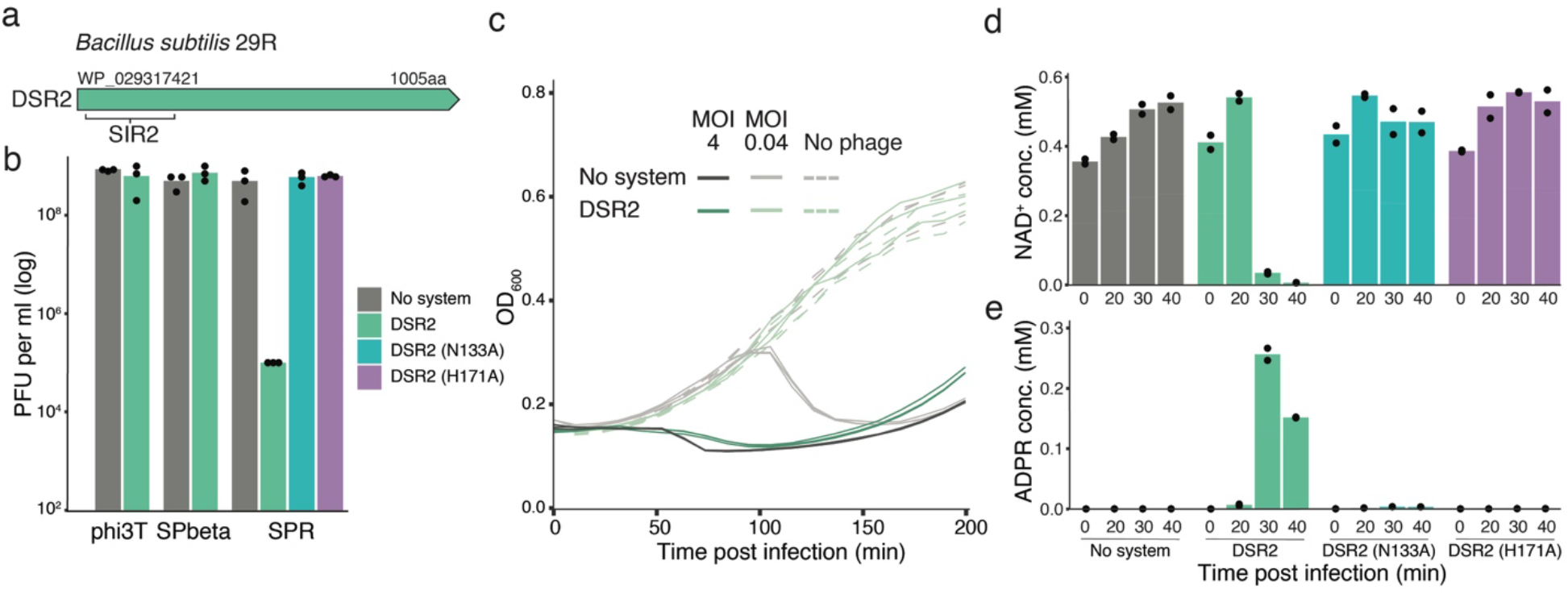
DSR2 is an abortive infection gene that causes NAD^+^ depletion in infected cells. (A) Domain organization of DSR2 from *B. subtilis* 29R. Protein accession in NCBI is indicated above the gene. (B) Efficiency of plating (EOP) for three phages infecting the control *B. subtilis* BEST7003 strain (no system) or *B. subtilis* BEST7003 with DSR2 cloned from *B. subtilis* 29R. For phage SPR, EOP is also presented for two mutations in the predicted SIR2 catalytic site. Data represent plaque-forming units (PFU) per milliliter. Bar graphs represent average of three independent replicates, with individual data points overlaid. (C) Liquid culture growth of DSR2-containing *B. subtilis* and control *B. subtilis* (no system), infected by phage SPR at 30 °C. Bacteria were infected at time 0 at an MOI of 4 or 0.04. Three independent replicates are shown for each MOI, and each curve represents an individual replicate. (D-E) Concentrations of NAD^+^ and ADPR in cell lysates extracted from SPR-infected cells as measured by targeted LC-MS with synthesized standards. X-axis represents minutes post infection, with zero representing non-infected cells. Cells were infected by phage SPR at a MOI of 5 at 30 °C. Bar graphs represent the average of two biological replicates, with individual data points overlaid. Colors are as in panel B.

We next tested whether DSR2 defends via abortive infection, a process that involves premature death or growth arrest of the infected cell, preventing phage replication and spread to nearby cells (Lopatina et al., 2020). When infected in liquid media, DSR2-containing bacterial cultures demised if the culture was infected by phage in high multiplicity of infection (MOI), similar to DSR2-lacking cells (Figure 1C). In low MOI infection, DSR2-lacking control cultures collapsed but DSR2-containing bacteria survived (Figure 1C). This phenotype is a hallmark of abortive infection, in which infected bacteria that contain the defense system do not survive but also do not produce phage progeny (Lopatina et al., 2020).

To ask whether DSR2 manipulates the NAD^+^ content of the cell during phage infection, we used mass spectrometry to monitor NAD^+^ levels in infected cells at various time points following initial infection. When DSR2-containing cells were infected by phage SPR, cellular NAD^+^ decreased sharply between 20 and 30 minutes from the onset of infection (Figure 1D). NAD^+^ levels did not change in cells in which the SIR2 active site was mutated, or in DSR2-lacking control cells, suggesting that the SIR2 domain is responsible for the observed NAD^+^ depletion (Figure 1D). In parallel with NAD^+^ depletion, we observed accumulation of the product of NAD^+^ cleavage, ADP-ribose (ADPR) (Figure 1E). These results demonstrate that DSR2 is an abortive infection protein that causes NAD^+^ depletion in infected cells.

DSR2 strongly protected *B. subtilis* cells against SPR, a phage belonging to the SPbeta group of phages (Kohm and Hertel, 2021) (Figure 1B). However, the defense gene failed to protect against phages phi3T and SPbeta, although both these phages belong to the same phage group as SPR (Figure 1B). This observation suggests that phages phi3T and SPbeta either encode genes that inhibit DSR2, or lack genes that are recognized by DSR2 and trigger its NADase activity. The phages SPR, phi3T and SPbeta all share substantial genomic regions with high sequence homology (Figure 2A). We therefore reasoned that co-infecting cells with two phages, either SPR and phi3T, or SPR and SPbeta, may result in recombination-mediated genetic exchange between the phages, which would enable pinpointing genes that allow escape from DSR2 defense when acquired by SPR (Figure 2B).

**Figure 2.**
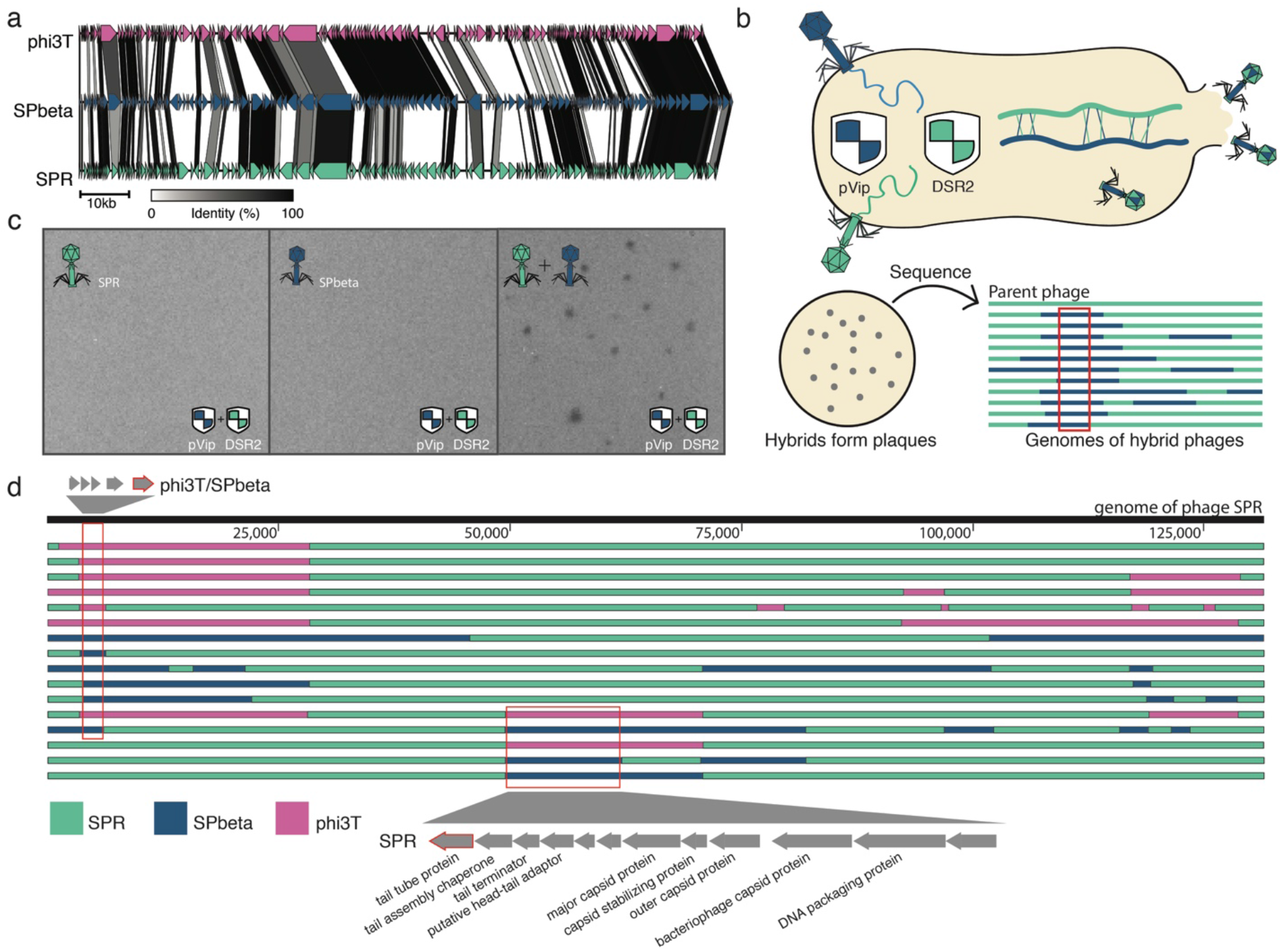
Genetic exchange between phages reveal regions responsible for escape from DSR2. (A) Genome comparison of phages SPR, SPbeta and phi3T was performed using clinker (Gilchrist and Chooi, 2021). Grey/black bands connect homologous genes, with shades of grey representing the % of amino acid sequence identity. (B) Schematic representation of the phage mating experiment. *B. subtilis* BEST7003 cells expressing both DSR2 and pVip are co-infected with phage SPR and SPbeta (or phi3T). Recombination between co-infecting phage genomes leads to hybrid phages that can overcome both defense systems and generate plaques. Examination of the genomes of multiple hybrids predicts genomic regions necessary to overcome defense. (C) Plaque assays with either one or two co-infecting phages. Cells expressing both DSR2 and pVip are infected either with phage SPR (left), phage SPbeta (middle) or both phages (right). (D) Hybrid phage genomes. Each horizontal line represents the genome of a hybrid phage that can overcome DSR2. Green areas are from phage SPR, blue from SPbeta and purple from phi3T. Representative non-redundant hybrid sequences are presented out of 32 sequenced hybrids. Red rectangles outline two areas that are predicted to allow the phage to overcome DSR2 defense. Top zoom inset shows genes found in the region acquired from phi3T or SPbeta, gene outlined in red codes for DSAD1. Bottom zoom inset shows genes present in the original SPR genome, outlined gene is tail tube protein.

To generate a bacterial host that would select for such genetic exchange events, we cloned into DSR2-containing cells an additional defense protein, a prokaryotic viperin homolog (pVip) from *Fibrobacter* sp. UWT3 (Bernheim et al., 2021). The pVip protein, when expressed alone in *B. subtilis*, protected it from phi3T and SPbeta but not from SPR, a defense profile which is opposite to that of the DSR2 profile in terms of the affected phages (Figure S1). Accordingly, none of the three SPbeta-group phages could form plaques on the strain that expressed both defensive genes. However, when simultaneously infecting the double defense strain with SPR and either phi3T or SPbeta, plaque forming hybrids were readily obtained, indicating that these hybrid phages recombined and acquired a combination of genes enabling them to escape both DSR2 and pVip (Figure 2C).

We isolated and sequenced 32 such hybrid phages, assembled their genomes, and compared these genomes to the genome of the parent SPR phage (Figure 2D; Supplementary File 1). This led to the identification of two genomic segments that were repeatedly acquired by SPR phages. Acquisition of either of these segments from a genome of a co-infecting phage rendered SPR resistant to DSR2 (Figure 2D). The first segment included five small genes of unknown function that are present in phages phi3T and SPbeta but not in the wild type SPR phage, and we therefore tested whether any of these genes was capable of inhibiting DSR2. One of the genes, when coexpressed with DSR2, rendered DSR2 inactive, suggesting that this phage gene encodes an anti-DSR2 protein (Figure 3A). This protein, which we named DSAD1 (DSR Anti Defense 1), is 120aa long and has no identifiable sequence homology to proteins of known function. Co-expression of DSR2 and tagged DSAD1, followed by pulldown assays, showed direct interaction between the two proteins (Figure 3B). These results indicate that DSAD1 is a phage protein that binds and inhibits DSR2.

**Figure 3.**
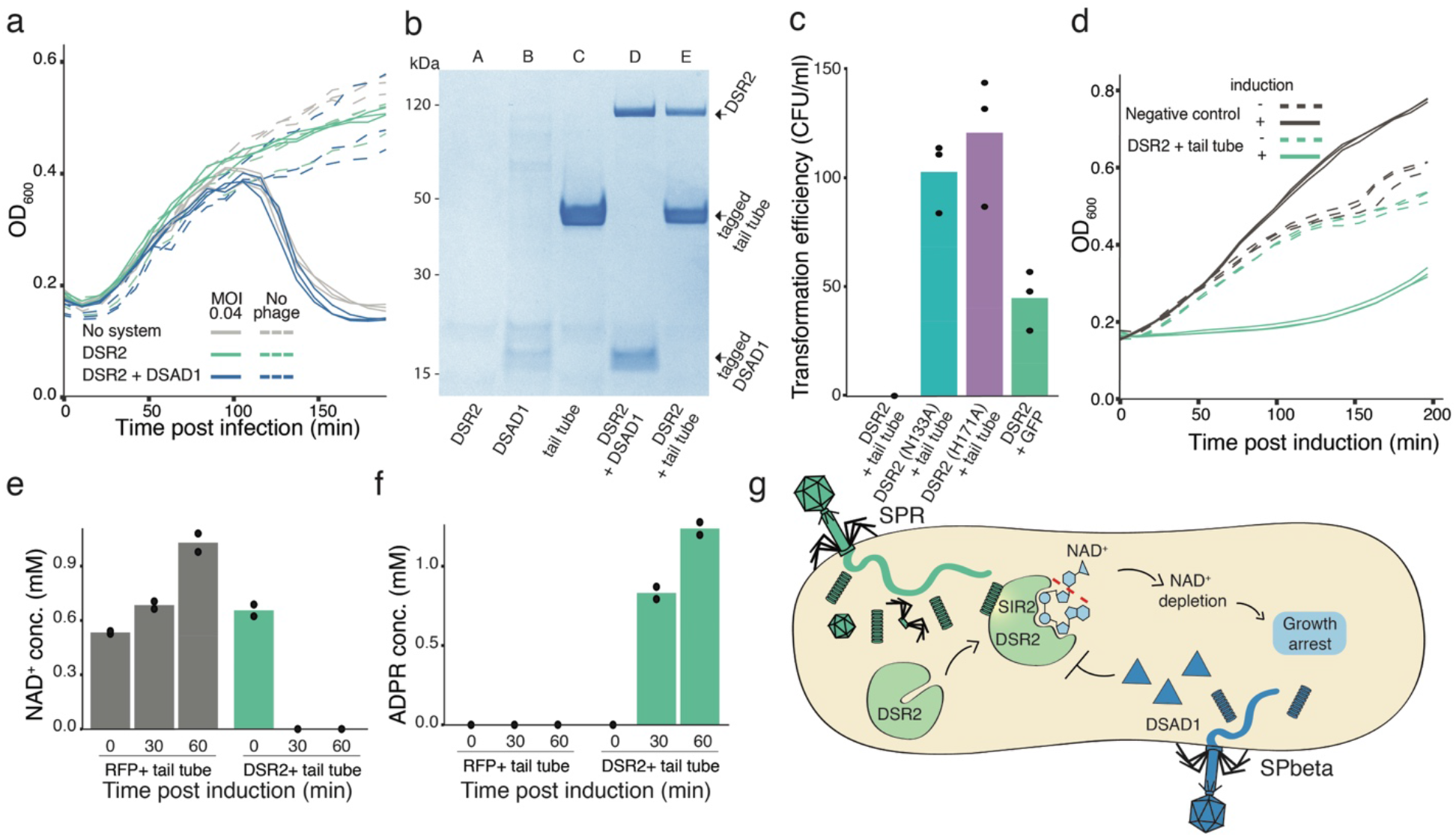
Phage proteins that activate and inhibit DSR2. (A) DSAD1 inhibits DSR2 defense. Liquid culture growth of *B. subtilis* BEST7003 cells expressing either DSR2 alone, or DSR2 and DSAD1, or control cells expressing neither gene, infected by phage SPR at 30 °C. Bacteria were infected at time 0 at an MOI of 0.04. Three independent replicates are shown for each MOI, and each curve shows an individual replicate. (B) Pulldown assays of the DSR2-DSAD1 and DSR2-tail tube complexes. DSAD1 and the tail tube protein of phage SPR were C-terminally tagged with TwinStrep tag. DSR2 in this experiment was mutated (H171A) to avoid toxicity. Shown is an SDS-PAGE gel. Lane A, cells expressing DSR2. Lane B, cells expressing tagged DSAD1. Lane C, cells expressing tagged tail tube protein from phage SPR. Lane D, cells co-expressing DSR2 and tagged DSAD1. Lane E, cells co-expressing DSR2 and tagged tail tube protein from SPR. (C) Transformation efficiencies of a vector containing the SPR tail tube protein or GFP control were measured for cells containing either WT DSR2 or two inactive DSR2 mutants. Y-axis represents the number of colony-forming units per milliliter. Bar graphs represent average of three replicates, with individual data points overlaid. (D) Liquid culture growth of *E. coli* that contains DSR2 and the tail tube gene of phage SPR, each under the control of an inducible promoter, and control *E. coli* that contains inducible GFP and DSR2 genes. Expression was induced at time 0 by adding 0.2% arabinose (for the tail tube gene) and 1 mM IPTG (for DSR2). Three independent replicates are shown, and each curve shows an individual replicate. (E-F) Concentrations of NAD^+^ and ADPR in cell lysates extracted from *E. coli* co-expressing DSR2 and SPR tail tube. X-axis represents minutes post expression induction, with zero representing non-induced cells. Control cells in this experiment express RFP and DSR2. Bar graphs represent the average of two biological replicates, with individual data points overlaid. (G) A model for the mechanism of action of DSR2. Phage infection is sensed by the recognition of the phage tail tube protein through direct binding to DSR2. This triggers the enzymatic activity of the SIR2 domain to deplete the cell of NAD^+^ thereby causing abortive infection. The phage anti-DSR2 protein DSAD1 inhibits DSR2 by direct binding.

We next examined the second genomic segment that, when acquired, allowed phage hybrids to escape DSR2. In the parent SPR phage, this region spans a set of genes encoding phage structural proteins, including capsid and tail proteins. Hybrid phages in which the original genes were replaced by their homologs from SPbeta or phi3T become resistant to DSR2 (Figure 2D). We hypothesized that one of the structural proteins in phage SPR is recognized by DSR2 to activate its defense, and when this protein is replaced by its homolog from another phage, recognition no longer occurs. To test this hypothesis, we attempted to clone each of the structural genes found in the original DNA segment in SPR into *B. subtilis* cells that also express DSR2. One of these genes, encoding a tail tube protein, could not be cloned into DSR2-expressing cells but was readily cloned into cells in which the DSR2 active site was mutated (Figure 3C).

To further test if co-expression of DSR2 and the SPR tail tube protein is toxic to bacteria, we cloned each of these genes under an inducible promoter within *E. coli* cells. In support of our hypothesis, growth was rapidly arrested in cells in which the expression of both genes was induced (Figure 3D), and these cells became depleted of NAD^+^ (Figure 3E, 3F). Furthermore, pulldown assays with tagged proteins showed that DSR2 directly binds the tail tube protein of phage SPR (Figure 3B). These results demonstrate that the tail tube protein of phage SPR is directly recognized by the defense protein DSR2, and that this recognition triggers the NADase activity of DSR2 and results in growth arrest (Figure 3G).

The original tail tube protein of phage SPR is substantially divergent from its counterparts from phages SPbeta or phi3T, sharing only ~40% amino acid sequence identity with these proteins (Figure S2). This divergence explains why the replacement of the original SPR protein with its SPbeta homolog renders the hybrid phage resistant to DSR2. Presumably, the evolutionary pressure imposed by DSR2 and other bacterial defense systems that recognize tail tube proteins has led to the observed diversification in these proteins in phages of the SPbeta group.

Our results show that DSR2 exerts its defensive activity by depleting NAD^+^ from infected cells. NADase activity was also recently described in the Thoeris defense system, in which a small molecule signal activates a SIR2-encoding effector to deplete NAD^+^ once phage infection has been recognized (Ofir et al., 2021). To test if NAD^+^ depletion is a general activity of SIR2 domains in bacterial defense systems, we examined three additional defense systems that contain a SIR2 domain (Figure 4A). These systems included a two-gene system that encodes, in addition to the SIR2 domain, also a prokaryotic argonaute homolog (pAgo), a two-gene system that encodes a HerA-like DNA translocase (Gao et al., 2020), and a single SIR2-domain protein called DSR1 (Gao et al., 2020) (Figure 4A). DSR1 was cloned into *B. subtilis* BEST7003, while the other two systems were cloned into an *E. coli* host. Consistent with our hypothesis, these systems defended against multiple different phages (Figure 4B, Figure S3), and NAD^+^ depletion was observed in each case (Figure 4C-E). Liquid infection with high and low MOIs showed a phenotype consistent with abortive infection for the SIR2/pAgo and the SIR2-HerA systems (Figure 4F-G). However, the DSR1 protein seems to inhibit the replication of phage phi29 without arresting the growth of the bacterial cells, and depletion of NAD^+^ was only transient (Figure 4E, 4H). Together, these results demonstrate a general role of SIR2 domains as NAD^+^ depleting effectors in bacterial defense against phage.

**Figure 4.**
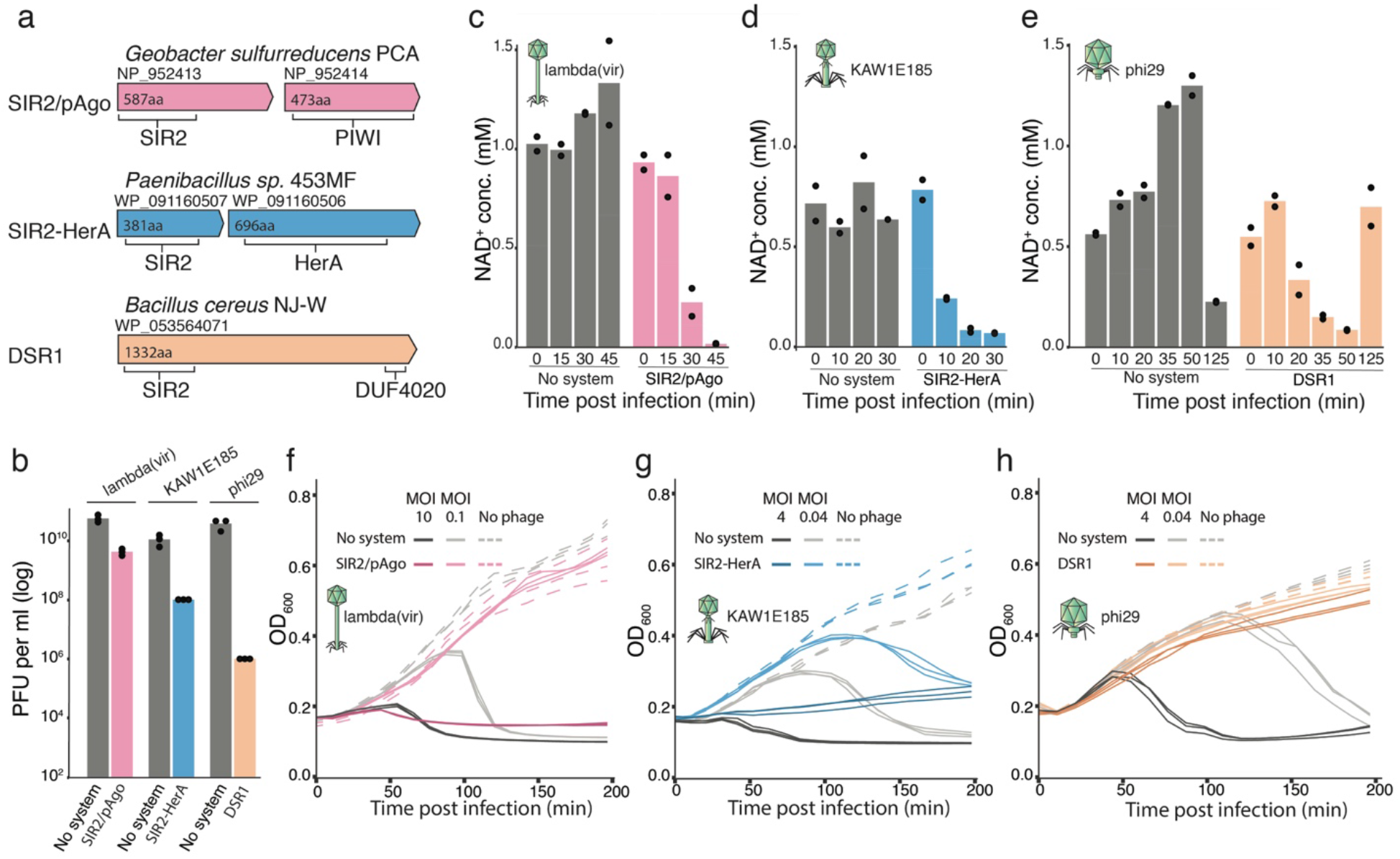
SIR2-containing defense systems deplete NAD^+^ in infected cells. (A) Domain organization of three defense systems that contain SIR2 domains. Protein accessions in NCBI are indicated. (B) Efficiency of plating for phages infecting defense-system-containing strains and control strains. SIR2-HerA and SIR2/pAgo were cloned into *E. coli* MG 1655, and DSR1 was cloned into *B. subtilis* BEST7003. Bar graphs are average of three biological replicates, with individual data points overlaid. KAW1E185 is short for vB_EcoM-KAW1E185, a T4-like phage. (C-E) Concentrations of NAD^+^ in cell lysates extracted from infected cells as measured by targeted LC-MS with synthesized standards. X-axis represents minutes post infection, with zero representing non-infected cells. “No system” are control cells that contain an empty vector instead of the defense system. Bar graphs represent the average of two biological replicates, with individual data points overlaid. (F-H). Liquid culture growth of bacteria that contain the respective defense system and control bacteria that contain an empty vector (no system). Bacteria were infected at time 0 at low or high MOIs, as indicated. Three independent replicates are shown for each MOI, and each curve shows an individual replicate.

## Discussion

Our data suggest that NAD^+^ depletion is a canonical function for SIR2 domains within bacterial defense systems. We show that four anti-phage defense systems, all containing SIR2 domains but otherwise comprising different architectures, deplete NAD^+^ in response to phage infection. Specifically, in the case of DSR2, we show that this protein recognizes newly translated phage tail tube proteins to become an active NADase. Phages of the SPbeta family have at least two versions of the tail tube protein, and only certain alleles are recognized by DSR2. In addition, we found that some phages in this family carry a small protein, DSAD1, which binds and inactivates DSR2.

NAD^+^ depletion was previously shown to be toxic to bacterial cells (Freire et al., 2019; Morehouse et al., 2020; Skjerning et al., 2019; Tang et al., 2018), and it was recently demonstrated in the Thoeris, CBASS and Pycsar systems that defense involving NAD^+^ depletion is associated with an eventual cell death (Morehouse et al., 2020; Ofir et al., 2021; Tal et al., 2021). Indeed, in three out of the four SIR2-containing systems that we studied, growth arrest or death of the bacterial host was observed in response to phage infection. However, DSR1 protection from phage phi29 did not involve culture collapse (Figure 4H). Furthermore, our data show that the NAD^+^ levels in cells containing DSR1 recovered after the initial depletion (Figure 4E). It is possible that in some cases, reversible reduction of NAD^+^ to low but not zero levels may be enough to interfere with phage replication while still allowing cell growth. Alternatively, lowered levels of NAD^+^ might have different outcomes for the infected cell depending on additional components derived from the infecting phage.

The arsenal of defense mechanisms known to protect bacteria against phage has recently been substantially expanded following the discovery of dozens of new anti-phage defense systems (Doron et al., 2018; Gao et al., 2020). While the mechanism of defense was deciphered for some of these systems (Bernheim et al., 2021; Cohen et al., 2019; Ofir et al., 2021; Tal et al., 2021), in many cases it is not known what molecular patterns of the phage trigger these systems to become active. In the current study we used a “phage mating” technique, promoting genetic exchange between phages that are sensitive to the defense system and phages that can overcome it. Genome analyses of hybrid phages enabled us to pinpoint the exact phage proteins that activate or repress DSR2. We believe that the phage mating approach should be a useful tool for other studies attempting to identify phage triggers of bacterial defense systems.

## Methods

### Bacterial strains and phages

*B. subtilis* strain BEST7003 (obtained from I. Mitsuhiro at Keio University, Japan) was grown in MMB (LB + 0.1 mM MnCl2 + 5 mM MgCl_2_, with or without 0.5% agar) at 30 °C. Whenever applicable, media were supplemented with spectinomycin (100 μg/ml) and chloramphenicol (5 μg/ml), to ensure selection of transformed and integrated cells. *E. coli* strain MG1655 (ATCC 47076) was grown in MMB at 37 °C. Whenever applicable, media were supplemented with ampicillin (100 μg/ml) or chloramphenicol (30 μg/ml) or kanamycin (50 μg/ml), to ensure the maintenance of plasmids. Phages used in this study are listed in Supplementary Table S1.

### Plasmid and strain construction

Details on defense systems analyzed in this study are summarized in Supplementary Table S2, and sequences of primers used in this study are in Supplementary Table S3. Defense systems DSR1, DSR2, and SIR2-HerA were synthesized by Genscript Corp. and cloned into the pSG1 plasmid (Doron et al., 2018) together with their native promoters. A whole operon of the SIR2/pAgo system, composed of the genes encoding the SIR2 (GSU1360, NP_952413.1) and pAgo (GSU1361, NP_952414.1) proteins, was amplified by PCR using the oligonucleotides MZ239 and MZ240, respectively, from the genomic DNA of *Geobacter sulfurreducens* Caccavo (LGC Standards cat #51573D-5). The resulting DNA fragment was digested by Eco31I (ThermoFisher cat #FD0293) and XhoI (ThermoFisher cat #FD0694) and using T4 DNA ligase (ThermoFisher cat #EL0014) was cloned into pBAD/HisA expression vector (ThermoFisher cat #V43001) precleaved with NheI (ThermoFisher cat #FD0973) and XhoI and dephosphorylated using FastAP (ThermoFisher cat #EF0651). The GsSir2 protein contains a His_6_-Tag at its N terminus. The mutants DSR2 (N133A) and DSR2 (H171A) were constructed using the Q5 Site-directed Mutagenesis kit (NEB, E0554S) using eaither primers JG216 and JG217, or JG220 and JG221 respectively.

A cloning shuttle vector for large fragments was constructed by Genscript Corp. This vector was constructed by replacing the Pxyl promoter and its downstream open reading frame in plasmid pGO1_thrC_Pxyl_cereus_ThsA (Ofir et al., 2021), with a synthesized Phspank sfGFP cassette taken from pDR111 (Overkamp et al., 2013), resulting in the plasmid pSG-thrC_Phspank_sfGFP (Supplementary File S2). The vector contains a p15a origin of replication and ampicillin resistance for plasmid propagation in *E. coli*, and a thrC integration cassette with chloramphenicol resistance for genomic integration into *B. subtilis*.

DSAD1 from SPbeta (NCBI protein accession WP_004399562) and phage tail tube protein from SPR (NCBI protein accession WP_010328117) were amplified from phage genomic DNA using primers JG346 and JG347 (for DSAD1) and JG142 and JG143 (for tail tube), and cloned into the pSG-thrC_Phspank_sfGFP vector, replacing sfGFP. The vector backbone was amplified using primers JG13 and JG14.

For expression in *E. coli MG1655*, DSR2 and DSR2(H171A) were amplified and cloned into the plasmid pBbA6c-RFP (Addgene, cat. #35290) using primers JG259 and JG260. The SPR tail tube gene was amplified from phage genomic DNA and cloned into the plasmid pBbS8k-RFP (Addgene, cat. #35276) with primers JG249 and JG250 for inducible expression in *E. coli*. For the assembly of tagged protein constructs, a pBbS8k with sfGFP fused to a C-terminal TwinStrep tag was first constructed (Supplementary File S3). The pBbS8k vector backbone was amplified using primers JG406 and JG407, and the sfGFP with a C-terminal TwinStrep tag was amplified from an *amyE* shuttle vector pGO6_amyE_hspank_GFP (Ofir et al., 2021) with primers JG366 and JG367 and cloned into the pBbS8k plasmid, replacing RFP. To create the C-terminally-tagged tail tube and DSAD1 constructs, the genes were amplified from phage genomic DNA using primers JG408 and JG409 (for tail tube) and JG410 and JG411 (for DSAD1) and cloned into the pBbS8k sfGFP with C-terminal TwinStrep mentioned above, replacing sfGFP. For this, the pBbS8k with C-terminal TwinStrep vector backbone was amplified using primers JG407 and JG412. All PCR reactions were performed using KAPA HiFi HotStart ReadyMix (Roche cat # KK2601). Cloning was performed using the NEBuilder HiFi DNA Assembly kit (NEB, E2621).

### *Bacillus* transformation

Transformation to *B. subtilis* BEST7003 was performed using MC medium as previously described (Wilson and Bott, 1968). MC medium was composed of 80 mM K_2_HPO_4_, 30 mM KH_2_PO_4_, 2% glucose, 30 mM trisodium citrate, 22 μg/ml ferric ammonium citrate, 0.1% casein hydrolysate (CAA), 0.2% sodium glutamate. From an overnight starter of bacteria, 10 μl were diluted in 1 ml of MC medium supplemented with 10 μl 1M MgSO_4_. After 3 hours of incubation (37°C, 200 rpm), 300 μl of the culture was transferred to a new 15 ml tube and ~200 ng of plasmid DNA was added. The tube was incubated for another 3 hours (37°C, 200 rpm), and the entire reaction was plated on Lysogeny Broth (LB) agar plates supplemented with 5 μg/ml chloramphenicol or 100 μg/ml spectinomycin and incubated overnight at 30°C.

### Plaque assays

Phages were propagated by picking a single phage plaque into a liquid culture of *B. subtilis* BEST7003 or *E. coli* MG1655 grown at 37°C to OD_600_ 0.3 in MMB medium until culture collapse. The culture was then centrifuged for 10 minutes at 3,200 x *g* and the supernatant was filtered through a 0.2 μm filter to get rid of remaining bacteria and bacterial debris. Lysate titer was determined using the small drop plaque assay method as described in Mazzocco et al. (2009).

Plaque assays were performed as previously described (Doron et al., 2018; Mazzocco et al., 2009). Bacteria containing defense system and control bacteria with no system were grown overnight at 37°C. Then 300 μl of the bacterial culture was mixed with 30 ml melted MMB 0.5% agar, poured on 10 cm square plates, and let to dry for 1 hour at room temperature. For cells that contained inducible constructs, the inducers were added to the agar before plates were poured. 10-fold serial dilutions in MMB were performed for each of the tested phages and 10 μl drops were put on the bacterial layer. After the drops had dried up, the plates were inverted and incubated overnight at room temperature or 37°C. Plaque forming units (PFUs) were determined by counting the derived plaques after overnight incubation and lysate titer was determined by calculating PFUs per ml. When no individual plaques could be identified, a faint lysis zone across the drop area was considered to be 10 plaques. Details of specific conditions used in plaque assays for each defense system are found in Supplementary Table S2.

### Liquid culture growth assays

Non-induced overnight cultures of bacteria containing defense system and bacteria with no system (negative control) were diluted 1:100 in MMB medium supplemented with appropriate antibiotics and incubated at 37°C while shaking at 200 rpm until early log phase (OD_600_ = 0.3). 180 μl of the culture were then transferred into wells in a 96-well plate containing 20 μl of phage lysate for a final MOI of 10 and 0.1 for phage lambda(vir), an MOI of 4 and 0.04 for phages SPR, phi29 and vB_EcoM-KAW1E185, or 20 μl of MMB for uninfected control. Infections were performed in triplicates from overnight cultures prepared from separate colonies. Plates were incubated at 30°C or 37°C (as indicated) with shaking in a TECAN Infinite200 plate reader and an OD_600_ measurement was taken every 10 min. Details of specific conditions used for each defense system are found in Supplementary Table S2.

### Liquid culture growth toxicity assays

Non-induced *E. coli* MG1655 with DSR2 and tail tube, or DSR2 and RFP (negative control), were grown overnight in MMB supplemented with 1% glucose and the appropriate antibiotics. Cells were diluted 1:100 in 3 ml of fresh MMB and grown at 37°C to an OD_600_ of 0.3 before expression was induced. The inducers were then added to a final concentration of 0.2% arabinose and 1mM IPTG, and MMB was added instead for the uninduced control cells. The cells were transferred into a 96-well plate. Plates were incubated at 37°C with shaking in a TECAN Infinite200 plate reader with an OD_600_ measurement was taken every 10 min.

### Cell lysate preparation for LC-MS

Overnight cultures of bacteria containing defense systems and bacteria with no system (negative control) were diluted 1:100 in 250 ml MMB and incubated at 37°C with shaking (200 rpm) until reaching OD_600_ of 0.3. For cells that contained inducible SIR2/pAgo constructs, the inducers were added at an OD_600_ of 0.1. A sample of 50 ml of uninfected culture (time 0) was then removed, and phage stock was added to the culture to reach an MOI of 5-10. Flasks were incubated at 30°C or 37°C (as indicated) with shaking (200 rpm) for the duration of the experiment. 50 ml samples were collected at various time points post infection. Immediately upon sample removal the sample tube was placed in ice, and centrifuged at 4°C for 5 minutes to pellet the cells. The supernatant was discarded and the tube was frozen at −80°C. To extract the metabolites, 600 μl of 100 mM phosphate buffer at pH 8, supplemented with 4 mg/ml lysozyme, was added to each pellet. Tubes were then incubated for 30 minutes at RT, and returned to ice. The thawed sample was transferred to a FastPrep Lysing Matrix B 2 ml tube (MP Biomedicals cat # 116911100) and lysed using FastPrep bead beater for 40 seconds at 6 m/s. Tubes were then centrifuged at 4°C for 15 minutes at 15,000 *g*. Supernatant was transferred to Amicon Ultra-0.5 Centrifugal Filter Unit 3 KDa (Merck Millipore cat # UFC500396) and centrifuged for 45 minutes at 4°C at 12,000 *g*. Filtrate was taken and used for LC-MS analysis. Details of specific conditions used for each defense system are found in Supplementary Table S2.

### Quantification of NAD^+^ and ADPR by HPLC-MS

Cell lysates were prepared as described above and analyzed by LC-MS/MS. Quantification of metabolites was carried out using an Acquity I-class UPLC system coupled to Xevo TQ-S triple quadrupole mass spectrometer (both Waters, US). The UPLC was performed using an Atlantis Premier BEH C18 AX column with the dimension of 2.1 × 100 mm and particle size of 1.7 μm (Waters). Mobile phase A was 20 mM ammonium formate at pH 3 and acetonitrile was mobile phase B. The flow rate was kept at 300 μl min^-1^ consisting of a 2 min hold at 2% B, followed by linear gradient increase to 100% B during 5 min. The column temperature was set at 25°C and an injection volume of 1 μl. An electrospray ionization interface was used as ionization source. Analysis was performed in positive ionization mode. Metabolites were detected using multiple-reaction monitoring, using argon as the collision gas. Quantification was made using standard curve in 0–1 mM concentration range. NAD+ (Sigma, N0632-1G) and ADPR (Sigma, A0752-25MG) were added to standards and samples as internal standard (0.5 μM). TargetLynx (Waters) was used for data analysis.

### Pulldown assays

Non-induced overnight cultures of *E. coli* MG1655 with DSR2 (H171A), DSAD1 with C-terminal TwinStrep tag, SPR tail tube protein with C-terminal TwinStrep tag, or combinations of these proteins were diluted 1:100 in 50 ml of MMB and grown at 37°C to an OD_600_ of 0.3. Expression was then induced by adding 0.2% arabinose and 1mM IPTG, and cells continued to grow to an OD_600_ of 0.9 at 37°C. Cells were centrifuged at 3,200 x *g* for 10 minutes. Supernatant was discarded and pellets were frozen in −80°C.

To pull down the proteins, 1 ml of Strep-Tactin wash buffer (IBA cat # 2-1003-100) supplemented with 4 mg/ml lysozyme was added to each pellet and incubated for 10 minutes at 37°C with shaking until thawed, and then resuspended. Tubes were then transferred to ice, and the resuspended cells transferred to a FastPrep Lysing Matrix B in 2 ml tube (MP Biomedicals cat # 116911100). Samples were lysed using FastPrep bead beater for 40 seconds at 6 m/s. Tubes were centrifuged for 15 minutes at 15,000 x g. Per each pellet, 30 μl of MagStrep “Type 3” XT beads (IBA cat # 2-4090-002) were washed twice in 300 μl wash buffer (IBA cat # 2-1003-100), and the lysed cell supernatant was mixed with the beads and incubated for 30-60 minutes, rotating at 4°C. The beads were then pelleted on a magnet, washed twice with wash buffer, and purified protein was eluted from the beads in 10 μl of BXT elution buffer (IBA cat # 2-1042-025). 30 μl of samples were mixed with 10 μl of 4X Bolt™ LDS Sample Buffer (ThermoFisher cat#B0008) and a final concentration of 1mM of DTT. Samples were incubated at 75°C for 5 minutes, and then loaded to a NuPAGE™ 4 to 12%, Bis-Tris, 1.0 mm, Mini Protein Gel, 12-well (ThermoFisher cat# NP0322PK2) in 20X Bolt™ MES SDS Running Buffer (ThermoFisher cat# B0002) and run at 160V. Gels were shaken with InstantBlue^®^ Coomassie Protein Stain (ISB1L) (ab119211) for 1 hour, followed by another hour in water. All bands shown in Figure 3B were verified to represent the indicated protein by mass spectrometry.

### Phage coinfection and hybrid isolation

50 μl overnight culture of *B. subtilis* containing DSR2 and pVip was mixed with 50 μl of phage SPR and 50 μl of either phi3T or SPbeta, each phage at a titer of 10^7^ PFU/ml. Bacteria and phages were left to rest at room temperature for 10 minutes before being mixed with 5ml of premelted MMB 0.3% agar and poured over a plate containing MMB 0.5% agar. Plates were left overnight at room temperature before being inspected for plaques. Single plaques were picked into 100 μl phage buffer (50 mM Tris-HCl pH 7.4, 100 mM MgCl_2_, 10 mM NaCl). Hybrid phages were tested for their ability to infect DSR2-containing cells using small drop plaque assay as described above.

### Sequencing and assembly of phage hybrids

High titer phage lysates (>10^7^ pfu/ml) of the ancestor and isolated phage hybrids were used for DNA extraction. 500 μl of the phage lysate was treated with DNaseI (Merck cat #11284932001) added to a final concentration of 20 μg/ml and incubated at 37°C for 1 hour to remove bacterial DNA. DNA was extracted using the QIAGEN DNeasy blood and tissue kit (cat #69504) starting from the Proteinase-K treatment step to lyse the phages. Libraries were prepared for Illumina sequencing using a modified Nextera protocol as previously described (Baym et al., 2015). Reads were de novo assembled using Spades3.14.0 (Prjibelski et al., 2020) with --careful flag set.

### Hybrid phage alignment

Hybrid phage genomes were aligned using SnapGene Version 5.3.2. Each hybrid genome was aligned to phage SPR and areas that did not align were aligned against the other phages in the coinfection experiment in order to verify their origin and gene content.

## Supporting information

File S1. Assembled genome sequences of 32 hybrid phages

File S2. Plasmid map of pSG-thrC_Phspank_sfGFP, shuttle vector for B. subtilis integration

File S3. Plasmid map of pBbS8k with C-terminal TwinStrep tag

## Acknowledgements

We thank the Sorek laboratory members for comments on the manuscript and fruitful discussion. We also thank Alexander Brandis and Tevie Mehlman from the Weizmann Life Sciences Core Facilities for targeted mass spectrometry analyses, and Alon Savidor from the Israel National Center for Personalized Medicine for protein mass spectrometry. R.S. was supported, in part, by the European Research Council (grant ERC-CoG 681203), Israel Science Foundation (grant ISF 296/21), the Ernest and Bonnie Beutler Research Program of Excellence in Genomic Medicine, the Deutsche Forschungsgemeinschaft (SPP 2330, grant 464312965), the Minerva Foundation with funding from the Federal German Ministry for Education and Research, and the Knell Family Center for Microbiology. V.S. was supported by the European Social Fund (grant 09.3.3-LMT-K-712-01-0126).

## Supplementary Materials

### Supplementary Figures

**Figure S1.**
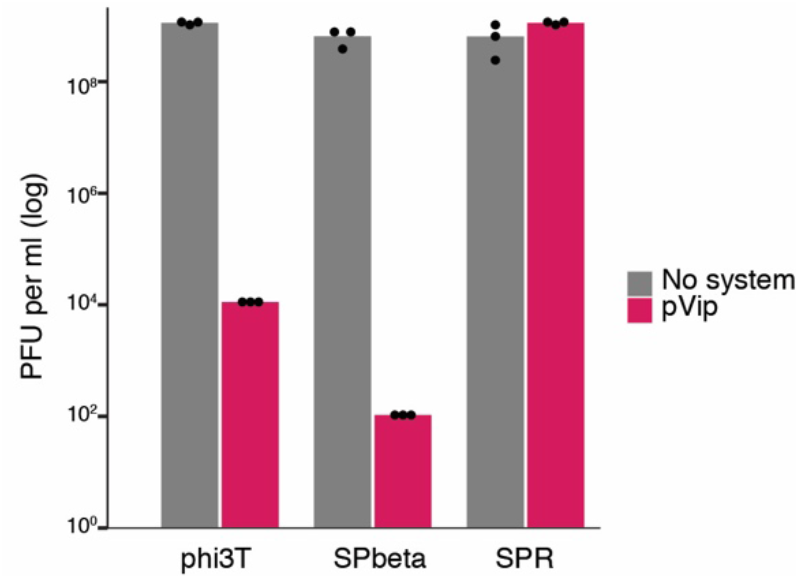
pVip alone protects against phi3T and SPbeta but not against SPR. Efficiency of plating (EOP) for three phages infecting the control *B. subtilis* strain (no system) or *B. subtilis* with pVip cloned from *Fibrobacter sp*. UWT3. Data represent plaque-forming units (PFU) per milliliter. Bar graphs represent average of three independent replicates, with individual data points overlaid. Control data are the same as those presented in Figure 1B.

**Figure S2.**
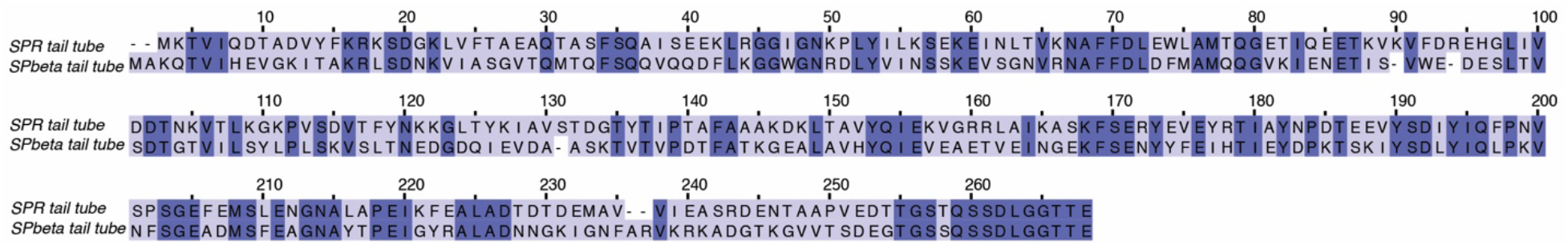
Multiple sequence alignment comparing the tail tube protein of SPR and SPbeta/phi3T. The NCBI accessions for the SPR and phi3T/SPbeta tail tube proteins are WP_010328117 and WP_004399252, respectively (the phi3T tail tube protein is identical to that of SPbeta). Protein amino acid sequences were aligned by Clustal Omega (Gabler et al., 2020).

**Figure S3.**
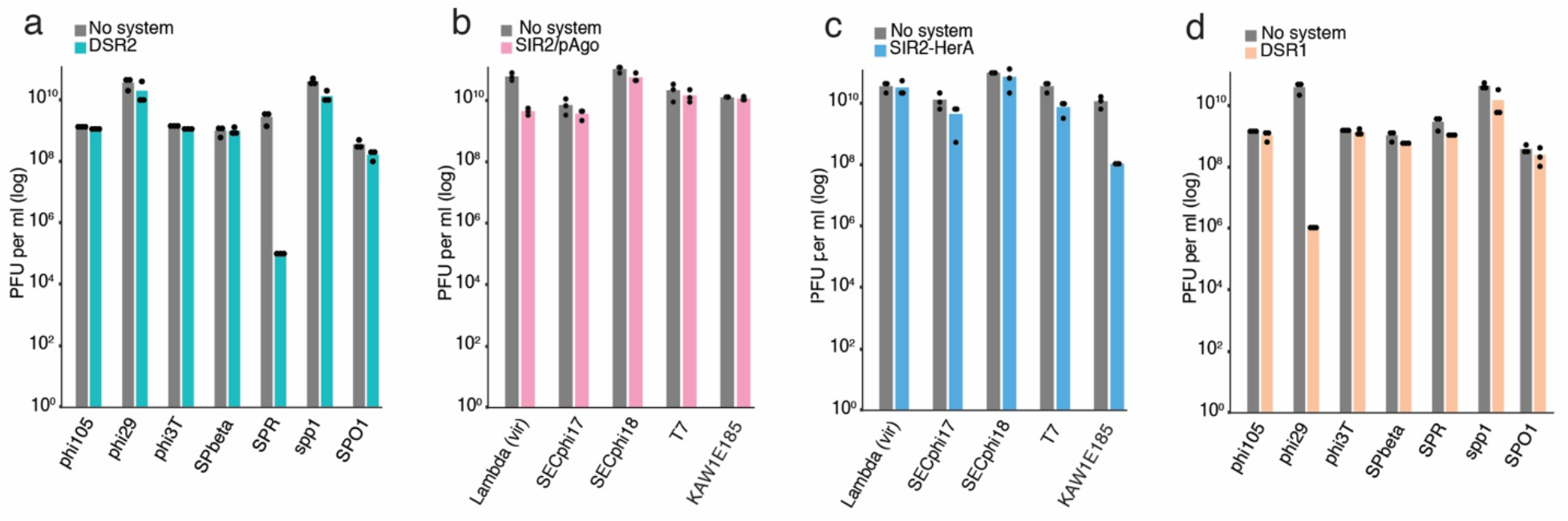
SIR2-containing defense systems protect against multiple phages. A-D. Efficiency of plating (EOP) for multiple phages infecting control bacteria (no system) or bacteria expressing defense systems. Data represent plaque-forming units (PFU) per milliliter. Bar graphs represent average of three independent replicates, with individual data points overlaid. KAW1E185 is short for vB_EcoM-KAW1E185, a T4-like phage. Data appearing in this figure also appear in Figure 4B. Data for control samples are the same in panels A and D.

### Supplementary Tables

**Table S1.** Phages used in this study.

| PHAGE | SOURCE | IDENTIFIER | ACCESSION |
| --- | --- | --- | --- |
| lambda(vir) | Udi Qimron |  | NC_001416.1 |
| phi105 | Bacillus Genetic Stock Center (BGSC) | BGSC (1L11) | HM072038.1 |
| phi29 | Deutsche Sammlung von Mikroorganismen und Zellkulturen (DSMZ) | DSM 5546 | NC_011048.1 |
| phi3T | Bacillus Genetic Stock Center (BGSC) | BGSC (1L1) | KY030782.1 |
| SECphi17 | Doron et al., 2018 |  | LT960607.1 |
| SECphi18 | Doron et al., 2018 |  | LT960609.1 |
| SPbeta | Bacillus Genetic Stock Center (BGSC) | BGSC (1L5) | AF020713.1 |
| SPO1 | Deutsche Sammlung von Mikroorganismen und Zellkulturen (DSMZ) | BGSC (1P4) | NC_011421.1 |
| Spp1 | Deutsche Sammlung von Mikroorganismen und Zellkulturen (DSMZ) | BGSC (1P7) | NC_004166.2 |
| SPR | Bacillus Genetic Stock Center (BGSC) | BGSC (1L56) |  |
| T7 | Udi Qimron |  | NC_001604.1 |
| vB_EcoM-KAW1E185 | Deutsche Sammlung von Mikroorganismen und Zellkulturen (DSMZ) | DSM 104099 | NC_054922.1 |

**Table S2.** SIR2 containing defense systems tested in this study.

| SYSTEM | HOST STRAIN | NEGATIVE CONTROL | PHAGES USED FOR MS AND LIQUID GROWTH ASSAYS | PHAGES USED FOR INFECTION PLAQUE ASSAYS | TEMP. (°C) | INDUCTION |
| --- | --- | --- | --- | --- | --- | --- |
| **DSR2** | *B. subtilis* | Empty cassette | SPR | phi29, spp1, SPR, SPbeta, phi3T, SPO1, phi105 | 30 | Native promoter |
| **DSR1** | *B. subtilis* | Empty cassette | phi29 | phi29, spp1, SPR, SPbeta, phi3T, SPO1, phi105 | 30 | Native promoter |
| **SIR2/PAGO** | *E. coli* | Empty pBAD | lambda(vir) | vB_EcoM-KAW1E185, lambda(vir), SECphi18, T7, SECphi17 | 37 | 0.2% arabinose |
| **SIR2-HERA** | *E. coli* | Empty vector | vB_EcoM-KAW1E185 | vB_EcoM-KAW1E185, lambda(vir), SECphi18, T7, SECphi17 | 37 | Native promoter |

**Table S3.**
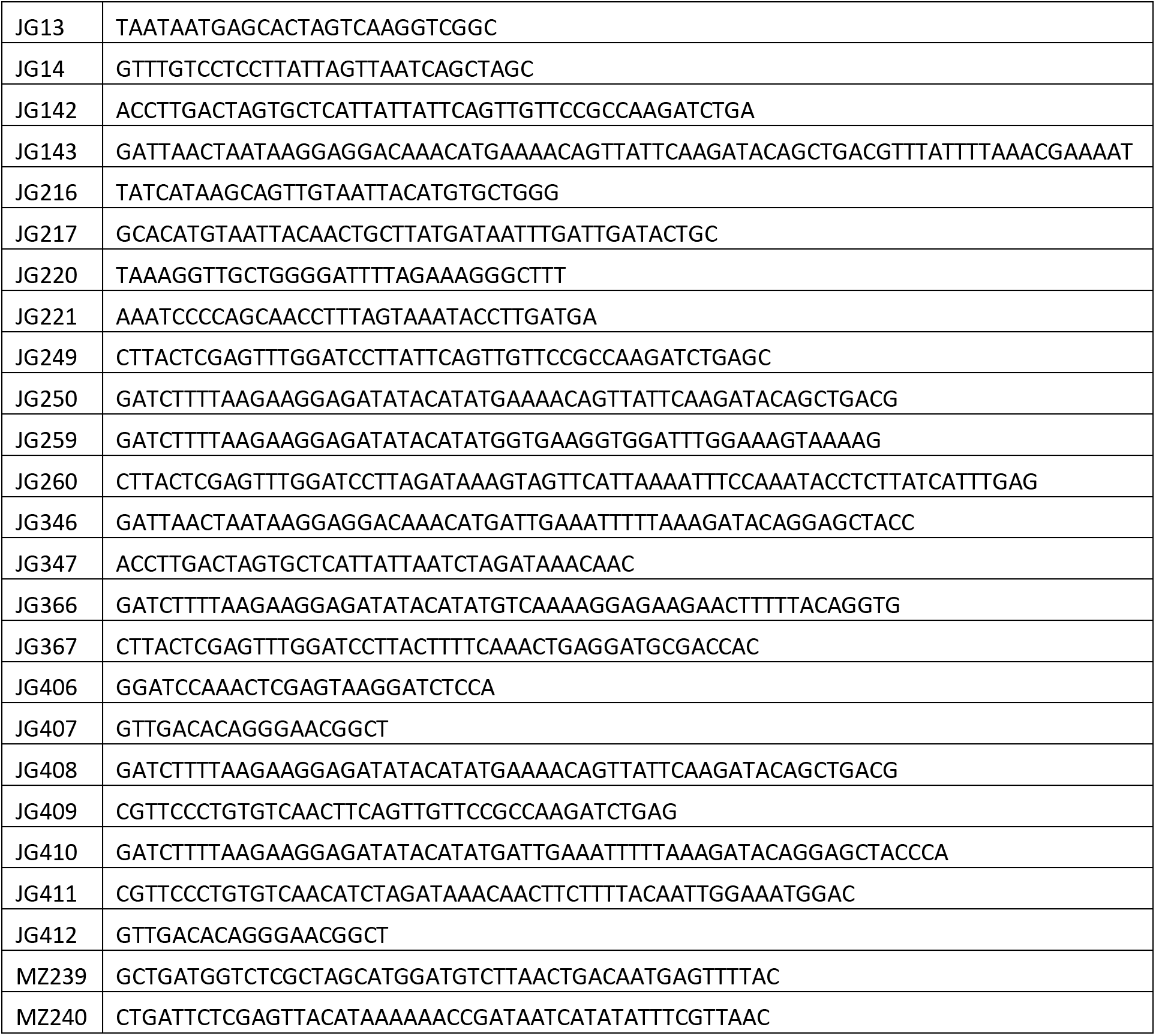
Primers used in this study.

### Supplementary Files

**File S1.** Assembled genome sequences of 32 hybrid phages

**File S2.** Plasmid map of pSG-thrC_Phspank_sfGFP, shuttle vector for *B. subtilis* integration

**File S3.** Plasmid map of pBbS8k with C-terminal TwinStrep tag

